# DeepVelo: Single-cell Transcriptomic Deep Velocity Field Learning with Neural Ordinary Differential Equations

**DOI:** 10.1101/2022.02.15.480564

**Authors:** Zhanlin Chen, William C. King, Aheyon Hwang, Mark Gerstein, Jing Zhang

## Abstract

Recent advances in single-cell RNA sequencing technology have provided unprecedented opportunities to simultaneously measure the gene expression profile and transcriptional velocity of individual cells, enabling us to sample gene regulatory network dynamics along developmental trajectories. However, traditional methods have faced challenges in modeling gene expression dynamics within individual cells due to sparse, non-linear (e.g., obligate heterodimer transcription factors), and high-dimensional measurements. Here, we present DeepVelo, a neural-network-based ordinary differential equation model that can learn non-linear, high-dimensional single-cell transcriptome dynamics and describe continuous gene expression changes within individual cells across time. We applied DeepVelo to multiple published datasets from different technical platforms and demonstrated its utility to 1) formulate transcriptome dynamics on different timescales, 2) measure the instability of cell states, and 3) identify developmental driver genes upstream of a signaling cascade. Benchmarking against state-of-the-art methods shows that DeepVelo can improve velocity field representation accuracy by at least 50% in out-of-sample cells. Further, perturbation studies revealed that single-cell dynamical systems may exhibit properties similar to those of chaotic systems. In summary, DeepVelo allows for the data-driven discovery of differential equations that delineate single-cell transcriptome dynamics.

**Teaser:** Embedding neural networks into ordinary differential equations to model gene expression changes within single cells across time.

## Introduction

Recent advances in single-cell RNA sequencing (scRNA-seq) have allowed gene expression profiling of individual cells, opening new avenues for investigating cellular development at an unprecedented resolution (*1*–*6*). A major goal in scRNA-seq is to study cell differentiation (*7*, *8*). In particular, understanding the regulatory cascade underlying cell state transitions is a crucial problem in embryonic development, tissue regeneration, and oncogenesis (*9*–*12*). However, due to the destructive nature of scRNA-seq, it is infeasible to track the transcriptomic profile of the same cell over time.

Several methods, e.g., *Monocle* and *Palantir*, have been proposed to model dynamic cell state transitions with pseudotime (*13*, *14*). Even though these methods have been successful in uncovering broad developmental trends, they have been limited in modeling the precise gene expression interactions driving cell transitions within individual cells (*15*). To estimate instantaneous cell transitions, RNA velocity used the ratio of spliced versus unspliced transcripts to infer the rate of change in the expression state with respect to time (*16*). While demonstrating encouraging results in various complex tissues, this method can only predict future cell states on the timescale of hours (*17*–*19*). To mitigate the short-term view of cell transitions, linear ordinary differential equations (ODEs) and sparse-approximation-based methods were proposed to predict the continuous evolution of cell states over long periods of time (*20*, *21*). However, linear models may fail to capture the non-linearity of gene expression dynamics. For example, obligate heterodimer transcription factors require the presence of both modules to activate the target gene, which is a known non-linear interaction that cannot be captured by linear models (*22*, *23*).

Inspired by recent developments in neural ODEs and data-driven dynamical systems (*24*, *25*), we present DeepVelo, a neural-network-based framework that can learn to formulate the dynamics underlying scRNA-seq experiments to overcome the aforementioned challenges. Our approach utilizes a denoising variational autoencoder (VAE) trained to predict the rate of change in gene expression (e.g., RNA velocity) from the gene expression state. We then embed the VAE into an ODE to model continuous changes in gene expression within an individual cell over time. Further, our deep-learning architecture can learn to remove noise from scRNA-seq measurements, reduce the dimensionality of the gene expression dynamics, and model complex non-linear gene interactions in a regulatory cascade.

Most importantly, by piecing together cells from different developmental stages, DeepVelo can learn to make cell state predictions farther into the future. In this regard, DeepVelo differs substantially from most single-cell methods, in that the objective of our framework is to derive neural-network-based differential equations describing continuous single-cell gene expression dynamics. To illustrate the robustness and general validity of our approach, we performed a proof-of-concept case study on mouse pancreatic endocrinogenesis (*26*). Then, we applied DeepVelo to decipher the gene expression dynamics behind the developing mouse brain, specifically in the dentate gyrus and neocortex (*27*, *28*). These samples represent scRNA-seq experiments from different tissues, technical platforms, and developmental timescales. With two additional data sources (mouse gastrulation, developing human forebrain), we further demonstrated the ability of DeepVelo to deconvolve gene co-expression networks and benchmarked our method against the state-of-the-art vector-field learning approach *SparseVCF* on out-of-sample velocity prediction accuracy (*16*, *26*, *29*).

## Results

### Neural ODEs for Modeling Single-cell Transcriptome Dynamics

In a gene regulatory network, the expression of certain genes can increase or decrease the expression of other genes. In a developmental context, a transitioning cell can signal a cascade of non-linear gene expression changes. One can formulate these gene-to-gene interactions as a function of time using differential equations. More specifically, each cell represents an instance of the dynamics sampled from the single-cell dynamical system. If the gene expression state of a cell is the vector 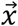, then the increase or decrease in gene expression with respect to time is the RNA velocity vector 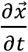. Rather than deriving a system of linear ODEs 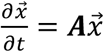 with matrix ***A***, we train a VAE 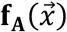 to learn the mapping from the gene expression state 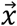 to the RNA velocity 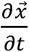 using data from each cell (Equation 1, Fig. 1a). Consequently, this VAE can model non-linear gene interactions when estimating the instantaneous gene expression change of individual cells. Then, given some initial gene expression state close to the data, we can numerically compute the future (or past) gene expression states with any black-box ODE solver. For example, given gene expression state 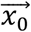 at time *t* = 0, we can use Euler’s method to find the gene expression state at 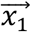 and can iteratively perform this step for 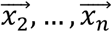 (Equation 2).

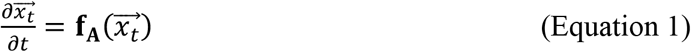

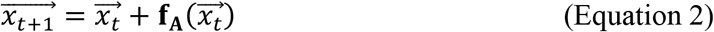

**Fig. 1.**
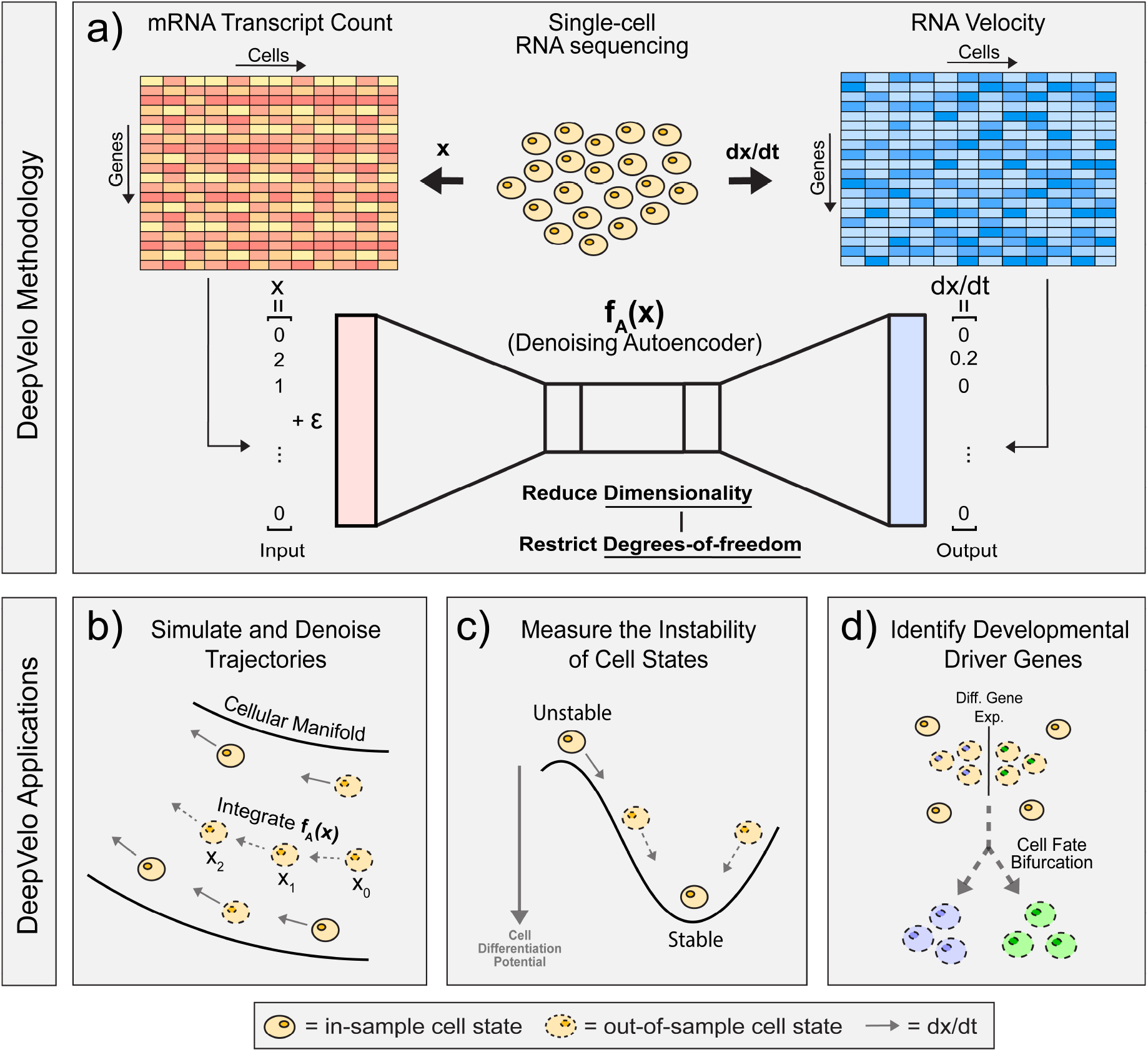
Schematic for DeepVelo. **a)** Gene expression profiles and corresponding transcriptional velocities can be derived from scRNA-seq data. After learning the mapping between gene expression and RNA velocity, the VAE represents a neural differential equation that encapsulates the transcriptome dynamics. **b)** Given an initial condition and time, our framework can solve for the future gene expression state by integrating the VAE with any black-box ODE solver. **c)** Our approach can simulate trajectories to evaluate the instability of cell states in a dynamical system. **d)** DeepVelo can perform *in silico* perturbation studies to identify the developmental driver genes that determine the fate of cell bifurcations

By sequentially computing the next gene expression state, DeepVelo can outline the developmental trajectory of single cells through time. Further, with different initial conditions 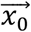, our framework can derive detailed insights into the future (or past) of different cell states. Here, we explored three applications of DeepVelo. First, we simulated and denoised developmental trajectories by extrapolating the dynamics to out-of-sample cells (Fig. 1b). Second, we evaluated the instability of cell states by tracking gene expression changes along simulated trajectories (Fig. 1c). Third, we performed *in silico* perturbation studies to investigate how initial gene expression conditions impact the fate of cell bifurcations (Fig. 1d.

### Predicting Future Cell States with Learned Neural ODEs

To evaluate whether DeepVelo can uncover dynamics from sparse and noisy scRNA-seq experiments, we considered a mouse pancreatic endocrinogenesis dataset with transcriptomes profiled at embryonic day E15.5 with the Chromium Single-cell 3’ Library from 10x Genomics (Fig. 2a) (*26*). Here, we show that summarizing the dynamics as neural ODEs can derive new insights about the data and further our understanding of pancreatic endocrinogenesis.

**Fig. 2.**
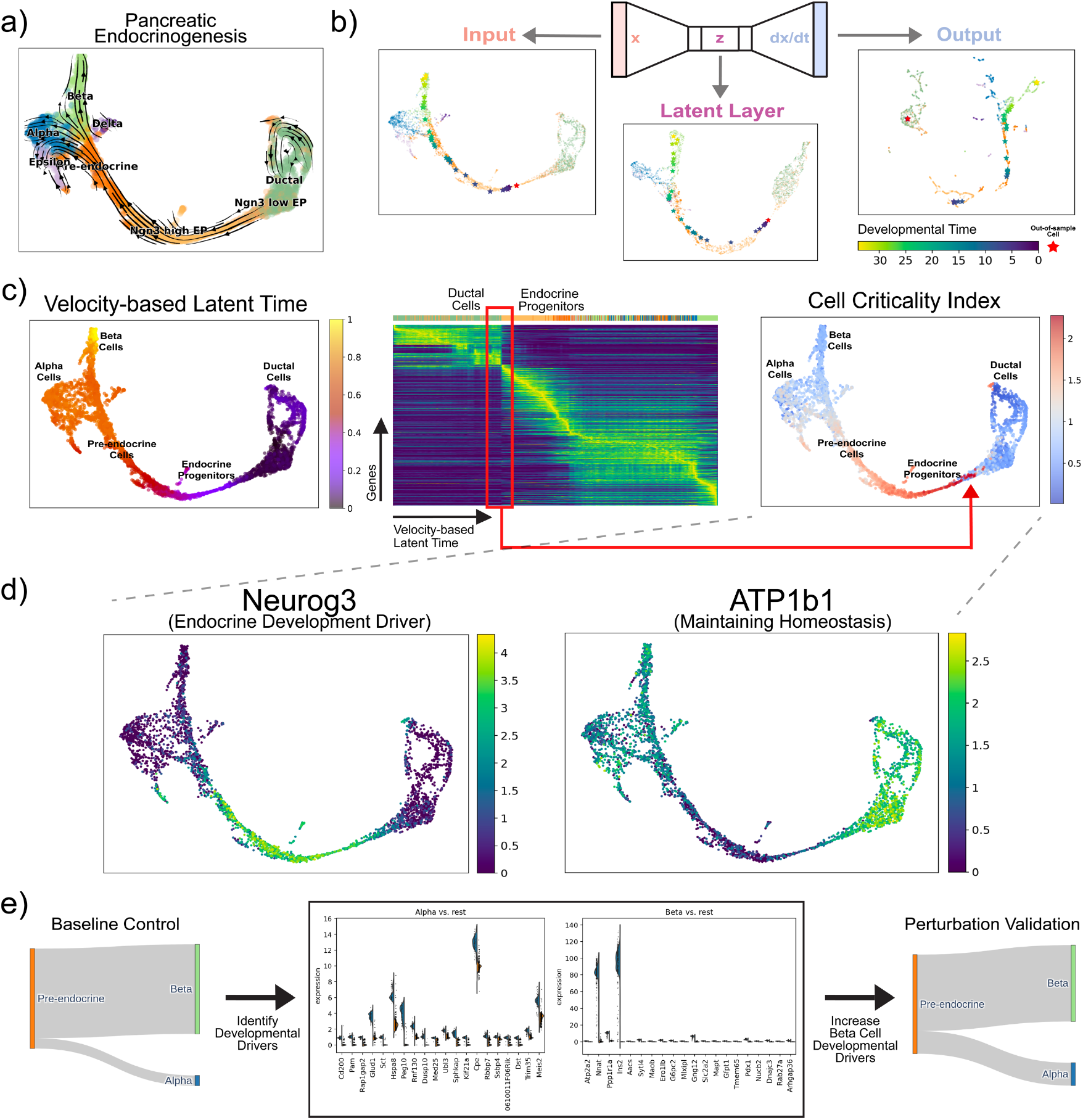
Pancreatic Endocrinogenesis as a Dynamical System. **a)** Pancreatic endocrinogenesis phase portraits projected in a low-dimensional embedding. Here, each cell is represented by a gene expression state and an RNA velocity vector. **b)** Simulating the developmental trajectory (in viridis) of an out-of-sample cell (in red) forward in time, visualized in uniform manifold approximation and projection (UMAP) embedding of the gene expression state (*x*), autoencoder latent layer (*z*), and velocity (*dx/dt*). **c)** Latent time derived from the RNA velocity for each cell (left). Gene expression of cells ordered by the velocity-derived latent time estimated by *scVelo* (middle). CCI derived from scDVF for each cell (right). Here, the CCI reveals unstable cell states indicative of fate commitment with a broad shift in gene expression patterns in latent time. **d)** Genes that highly correlate with the CCI reveal driving forces behind endocrine progenitor dynamics. In particular, *Neurog3*, which positively correlates with the CCI, is a known pre-endocrine master regulator, and *ATP1b1*, which negatively correlates with the CCI, has an important function in maintaining homeostasis in stable and stationary cell types. These genes support the CCI as a metric for the stability of single-cell states. **e)** Differential gene expression analysis of simulated pre-endocrine cells reveals key putative genes that correlate with the fate of transforming into an alpha versus a beta cell. Early perturbation of the top differentially expressed genes associated with an alpha cell fate resulted in a higher proportion of alpha cells from the perturbed pre-endocrine cells, suggesting a causal relationship through *in silico* studies

First, we examined a hypothetical trajectory simulated from DeepVelo after training the VAE on pancreatic endocrinogenesis cell states and velocities. When simulating hypothetical trajectories, future state predictions rely on out-of-sample cell states predicted from the previous time point. Hence, we evaluated the ability of DeepVelo to predict future cell states for an out-of-sample initial condition. We simulated an out-of-sample cell by adding noise to the gene expression state of an existing cell, thereby representing a cell state that did not previously exist in the data. The simulated developmental path shows that our predicted gene expression states moved along existing trajectories in the data manifold (Fig. 2b). In pancreatic endocrinogenesis, the out-of-sample cell started as an endocrine progenitor, developed into a pre-endocrine cell, and ultimately became a beta cell. When we solve the VAE with evenly distributed time increments, the distances between intermediate states reflect the magnitude of the RNA velocity vectors. Higher rates of change in gene expression generated more separated intermediate states. Conversely, lower rates of change produced a denser collection of intermediate points along the manifold.

During VAE training, regularized loss in the latent layer promotes a continuous, compact representation of locally similar cell states and trajectories in the embedding. When the VAE is trained to model gene expression dynamics, the latent layer integrates gene expression state and RNA velocity information to generate an embedding of the low-dimensional dynamic manifold (Fig. 2b). In pancreatic endocrinogenesis, the chronological and hierarchical orders of developmental trajectories are properly encoded in the latent layer, similar to gene expression embeddings. For example, ductal cells represent a major starting state whereas alpha and beta cells represent a major terminal state. The simulated cell migrates along existing trajectories in the low-dimensional dynamic manifold.

### Characterizing Instability in DeepVelo-predicted Trajectories with the Cell Criticality Index

Next, we aimed to characterize the stable and unstable fixed points of this single-cell dynamical system. By looking forward in time, we can numerically approximate the cell criticality index (CCI), which describes the instability of single-cell states. For a cell, we define the CCI as the cumulative information change, or the cumulative Kullback–Leibler (KL) divergence, between gene expression distributions at each time step in the developmental trajectory. In other words, cell states that undergo large changes across time will have a high CCI, whereas cell states that go through only small changes will have a low CCI.

For each cell, we used DeepVelo to compute a developmental path such that the cell arrived at a steady terminal state. Then, we calculated the CCI along each path (Fig. 2c). The resulting developmental topology is similar to the classical Waddington landscape (*30*). In particular, the CCI can reveal unique topological information in the developmental landscape not directly observed in the velocity-based latent time estimated by *scVelo* (*31*). For example, the endocrine progenitor states exhibit a higher criticality, whereas the ductal cell and differentiated endocrine cell states experience a lower criticality. When ductal cells undergo transformation into islet cell types, the heightened criticality in endocrine progenitors represents fate commitment or a point of no return during development. In dynamical systems, this behavior suggests that cell states with low criticality are located at a stable fixed point, with the cell identity remaining stable against small gene expression perturbations. More interestingly, the endocrine progenitors are located at an unstable fixed point with properties similar to those of a chaotic system in which a small perturbation may result in large downstream changes. We can substantiate the instability of cell states by examining the genes that best correlate with the CCI (Fig. 2d). For example, previous experiments have shown that *Neurog3*, which positively correlates with the CCI, is a known driver for endocrine commitment, and *ATP1b1*, which negatively correlates with the CCI, is important for maintaining homeostasis in stable and stationary cell types (*32*, *33*). The expression of these genes supports the CCI as a metric for evaluating the instability of single-cell states. Further, when viewed in conjunction with velocity-based latent time, the CCI can provide new insights into the relative timing of critical cell states in a developmental process.

### Conducting *in Silico* Perturbation Studies with DeepVelo

Lastly, we investigated the behavior of this dynamical system with similar perturbation studies pioneered by (*34*). The goal of *in silico* perturbation studies is to computationally identify which initial gene expression conditions impact the fate of cell bifurcations. In short, we randomly sampled pre-endocrine cells (*n* = 1,000) as the initial conditions. By allowing these simulated pre-endocrine cells to naturally evolve according to the dynamics learned by DeepVelo, we observed a baseline 9:1 ratio of terminal beta versus alpha cell states. The ratio of terminal cell states indicates that the beta cell state is a stronger attractive terminal state than the alpha cell state, which corroborates with previous conclusions (*31*). Then, we examined differential gene expression between initial conditions of different fates. The results suggest that early expression perturbations in key upstream genes correlate with the fate of developmental bifurcations (Fig. 2e).

### Exploring the Neural ODEs Governing the Developing Mouse Dentate Gyrus

Here, we evaluated whether DeepVelo can uncover the dynamics of a dataset from a different tissue, developmental timescale, and technical platform. We considered an scRNA-seq experiment of the developing mouse dentate gyrus with transcriptomes profiled using droplet-based scRNA-seq (Fig. 3a) (*27*). After obtaining a neural network representation of the dentate gyrus dynamics, we simulated an out-of-sample cell by perturbing the gene expression state of an *Nbl2* cell. With the out-of-sample cell as the initial condition, we used DeepVelo to simulate an out-of-sample cell trajectory, which moved along the existing granule cell trajectory in the data (Fig. 3b). Further, the VAE embeddings properly encoded the developmental hierarchy of cell types in the low-dimensional dynamic manifold (Fig. 3c).

**Fig. 3.**
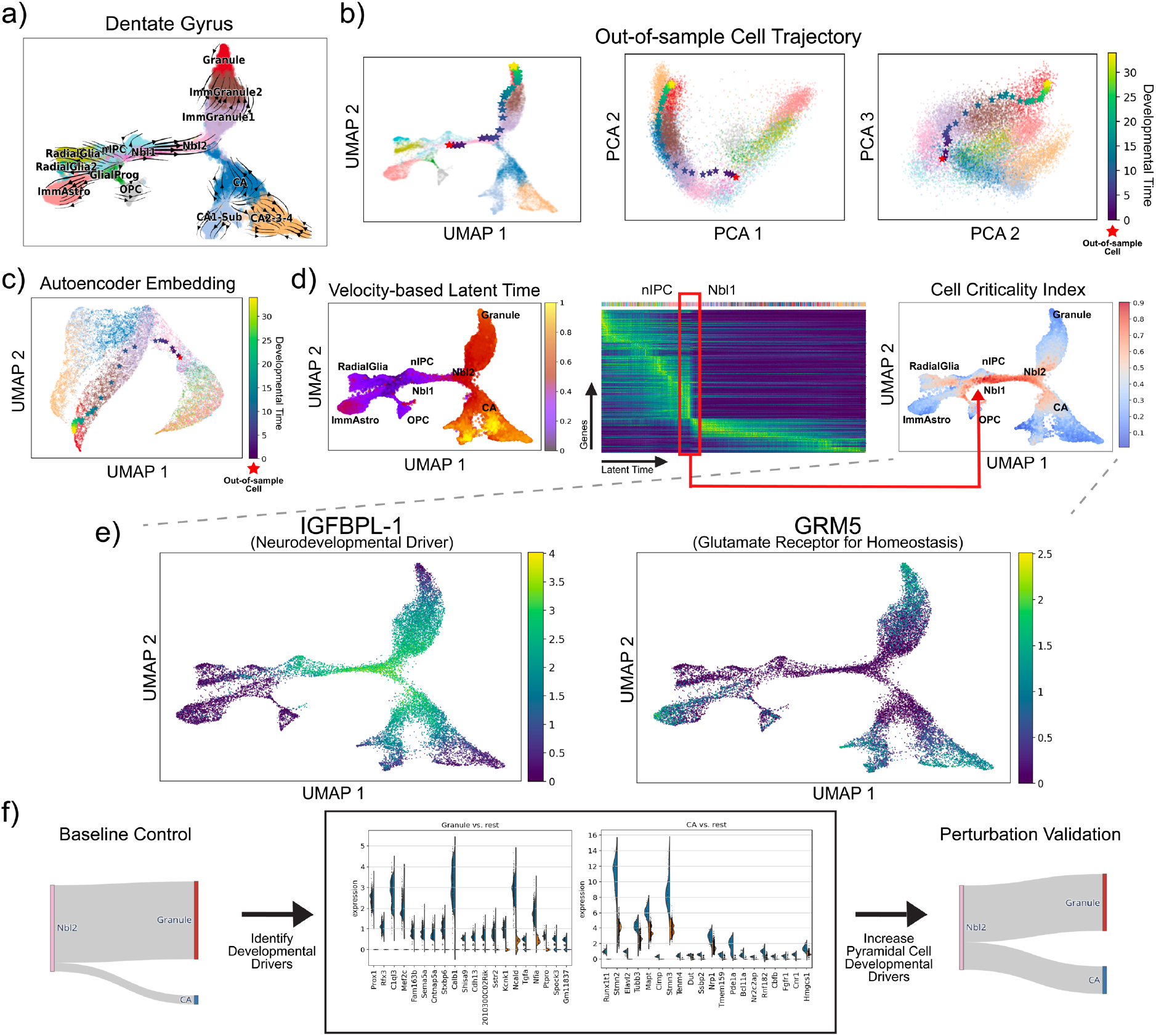
Developing Mouse Dentate Gyrus as a Dynamical System. **a)** Dentate gyrus phase portraits projected in a low-dimensional embedding. Here, each cell is represented by a gene expression state vector and an RNA velocity vector. **b)** Simulating the developmental trajectory (in viridis) of an out-of-sample cell (in red) forward in developmental time, visualized in UMAP and principal component analysis (PCA) embeddings. **c)** Visualizing the low-dimensional dynamic manifold of the high-degree-of-freedom single-cell developmental process, with the simulated out-of-sample cell trajectory (in red and viridis). **d)** Velocity-based latent time derived from *scVelo* for each cell (left). Gene expression of cells ordered by velocity-derived latent time (middle). CCI derived from scDVF for each cell (right). The CCI reveals instability during the transition from *nIPC* to *Nbl1* cells, indicative of cell fate commitment. **e)** Genes that highly correlate with the CCI reveal driving forces behind dentate gyrus dynamics. In particular, *IGFBPL-1*, which shows the strongest positive correlation with the CCI, regulates neurodevelopment, and *GRM5*, which shows the strongest negative correlation with the CCI, encodes glutamate receptors in stable and stationary neurons. These genes further substantiate the CCI as a metric for the stability of single-cell states. **f)** Differential gene expression analysis of simulated pre-endocrine cells reveals key putative genes that correlate with the fate of transforming into a pyramidal versus granule cell. Early perturbation of the top differentially expressed genes associated with a pyramidal cell fate resulted in a higher proportion of pyramidal cells from the perturbed *Nbl2* cells, suggesting a causal relationship through *in silico* studies.

When examining critical cell states in the dentate gyrus, we observed an abrupt gene expression shift in the developmental manifold, which can be visualized by ordering cells in latent time derived from *scVelo* (Fig. 3d). Specifically, the change in gene expression marks the transition from *nIPC* to *Nbl1* cells and suggests fate commitment during the transition. After calculating the CCI, we found that cells experiencing this abrupt change also have a high criticality, which substantiates the CCI as a metric for quantifying the instability of cell states. In addition, the most strongly correlated genes in the dentate gyrus highlight the robustness of the CCI as an instability measure. For example, *IGFBPL-1*, which shows the strongest positive correlation with the CCI, drives neuron differentiation in progenitor cells, and *GRM5*, which shows the strongest negative correlation with the CCI, encodes glutamate receptors in stable and differentiated neurons (Fig. 3e) (*35*, *36*).

Lastly, we conducted *in silico* perturbation studies to determine the genetic drivers governing dentate gyrus cell fate decisions. We randomly sampled upstream *Nbl2* cells (*n* = 1,000) as the initial conditions and allowed the simulated *Nbl2* cells to naturally evolve according to the dynamics captured by DeepVelo, which resulted in either terminal granule or pyramidal cell states. Then, we performed differential expression analysis on the initial conditions (i.e., the simulated *Nbl2* cell states) of different fates (Fig. 3f). The top differentially expressed gene associated with a granule cell fate was *Prox1*. This gene has been previously identified by RNA velocity and experimentally validated as being necessary for granule cell formation; moreover, the deletion of *Prox1* leads to adoption of the pyramidal neuron fate (*35*). In addition, DeepVelo identified the top pyramidal neuron developmental driver gene as *Runx1t1*, which was recently shown to induce pyramidal neuron formation, with its deletion resulting in reduced neuron differentiation *in vitro* (*37*). As further validation, we increased the expression of pyramidal neuron developmental driver genes in simulated *Nbl2* cells and observed an elevated proportion of pyramidal neurons as terminal states (from 10% to 30%; binomial test *p* < 10^-7^) under the dynamics captured by DeepVelo. In summary, *in silico* perturbation studies can provide a low-cost alternative for identifying developmental driver genes. Further, the results show that DeepVelo is robust for scRNA-seq datasets from different tissues, developmental timescales, and technical platforms.

### Formulating Neural ODEs Underlying the Mouse Neocortex Across Multiple Embryonic Days

We evaluated whether DeepVelo can learn the dynamics underlying a more complex developmental process. We examined an scRNA-seq experiment of the developing mouse neocortex with transcriptomes profiled at embryonic days E14, E15, E16, and E17 (*28*). After batch correction and velocity-field learning, we generated an out-of-sample intermediate progenitor (IP) as the initial condition. The simulated developmental trajectory suggests that the hypothetical IP transitioned into a migrating neuron (MN) and then subsequently became a CorticoFugal neuron (CFN). The distance between each time step was larger when the simulated cell was an IP compared with an MN, indicating that the IPs transition at a faster rate than the MNs (Fig. 4a). The latent layer embeddings also properly encoded the progression from neural progenitors to differentiated neurons (Fig. 4b).

**Fig. 4.**
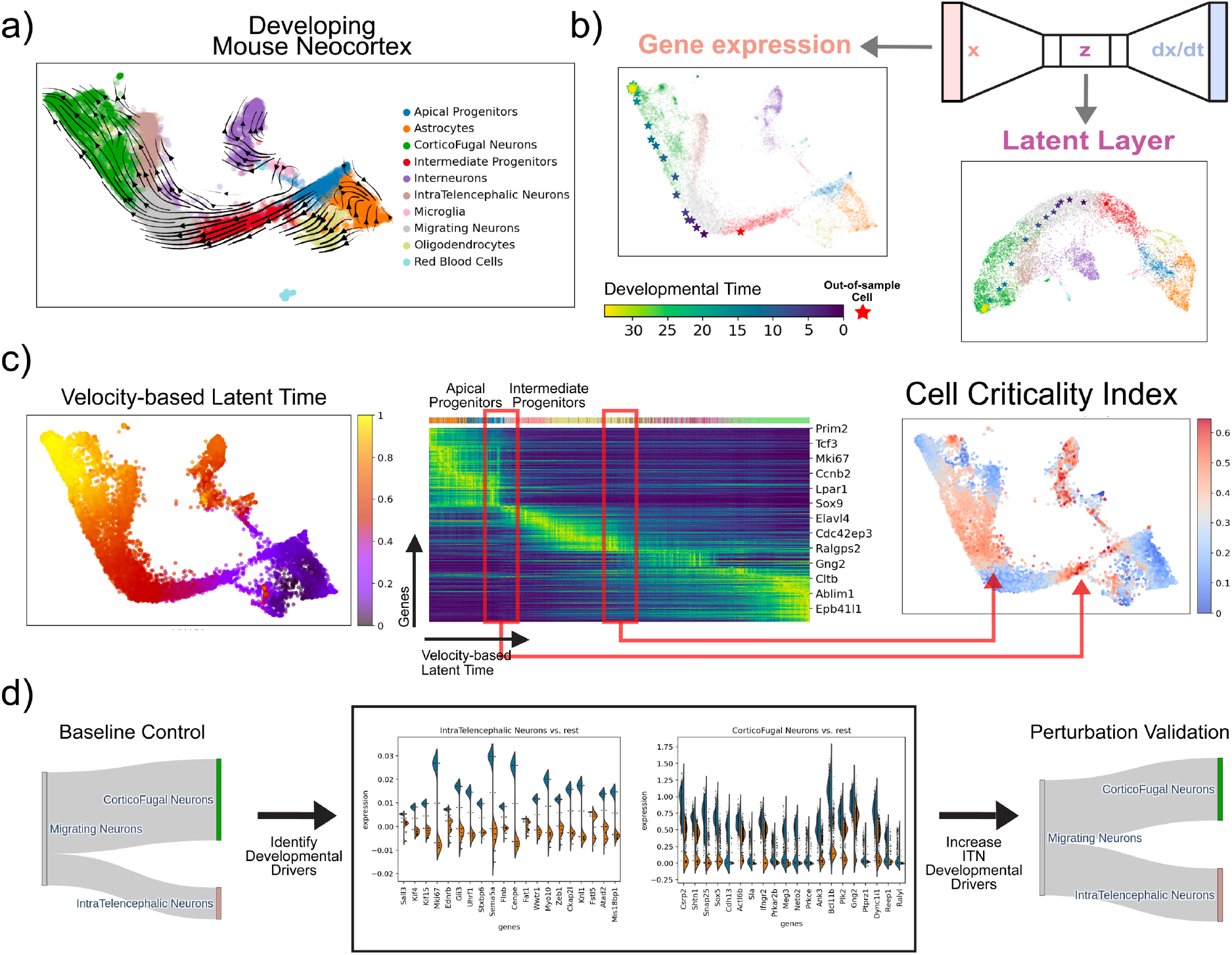
Developing Mouse Neocortex Over Multiple Embryonic Days. **a)** Mouse neocortex phase portraits projected in a low-dimensional embedding. **b)** Simulating the trajectory (in viridis) of an out-of-sample cell (in red) forward in developmental time, visualized in gene expression embeddings and VAE latent layer embeddings. **c)** The CCI reveals two unstable fixed points indicative of a biphasic fate-commitment dynamic. One point highlights the transition from APs to IPs, and the other highlights the transition from MNs to CFNs. **d)** Differential gene expression analysis of the simulated MNs reveals key putative genes that correlate with the fate of transforming into an ITN versus CFN. Early perturbation of the top differentially expressed genes associated with a pyramidal cell fate resulted in a 10% higher proportion of ITNs from the perturbed MNs.

When analyzed in conjunction with velocity-based latent time, we observed a biphasic fate-commitment dynamic with high criticality among IPs and MNs but low criticality in apical progenitors (APs) through the CCI (Fig. 4c). This finding suggests that APs play a more important role in self-renewal, whereas the IPs are pre-programmed for neuronal differentiation through a broader shift in gene expression patterns. Additionally, as MNs branch into IntraTelencephalic Neurons (ITNs) and CFNs, DeepVelo can effectively disentangle the underlying bifurcation dynamics (Fig. 4d). When we allowed a set of simulated MNs (*n* = 1000) to evolve according to the dynamics learned by DeepVelo, we observed a baseline 3:7 ratio of ITNs versus CFNs. After perturbing the initial differentially expressed genes associated with an ITN fate in another set of simulated MNs (*n* = 1000), we observed a 10% increase in ITN proportion compared with the baseline. These observations indicate that DeepVelo can accurately learn an overarching neural ODE of mouse neocortex gene expression dynamics spanning several embryonic days. Further, the results suggest that our framework can identify driver genes that play crucial roles in developmental dynamics with a continuous representation of velocity fields learned by DeepVelo.

### Reconstructing Gene Co-expression Networks with Retrograde Trajectories

RNA velocity can be used to predict gene expression changes in individual cells on the timescale of hours. Previous simulations in this study used hypothetical progenitor cells as the initial conditions and computed future trajectories resulting in differentiated cells as terminal states. Conversely, we can use differentiated (or terminal) cells as the initial conditions and reverse time with DeepVelo. In this case, the retrograde developmental trajectory represents the gene dynamics that would have resulted in the terminal cell types.

Due to sparse and noisy measurements, it is often challenging to detect strong correlations between genes in scRNA-seq, thereby making it difficult to find coherent functional modules in gene co-expression networks (*38*–*40*). However, denoising VAEs in DeepVelo can reduce the variability along a developmental trajectory caused by the sparsity and noise associated with scRNA-seq (Fig. 5a). We hypothesize that cells in denoised trajectories simulated from DeepVelo (with a representative initial condition) can amplify correlations within functional gene modules (Fig. 5b). Indeed, the gene co-expression network of beta cells in retrograde trajectories has more significant gene correlations than co-expression networks from beta cells measured by scRNA-seq. Further, we biclustered the co-expression matrix into gene clusters. By benchmarking our approach on four datasets, we demonstrated that the gene clusters discovered from our method are more coherent by comparing the gene ontology (GO) enrichments. The benchmarks show that functional gene modules found from denoised and dynamic cells in retrograde trajectories have an enrichment for cell-type-specific GO terms that is at least two orders of magnitude higher than that for static cell clusters (Fig. 5c). Therefore, the retrograde trajectories computed by DeepVelo can effectively disentangle trajectory-specific gene regulatory networks and can provide a computational solution for boosting signal-to-noise ratios in single-cell gene co-expression networks.

**Fig. 5.**
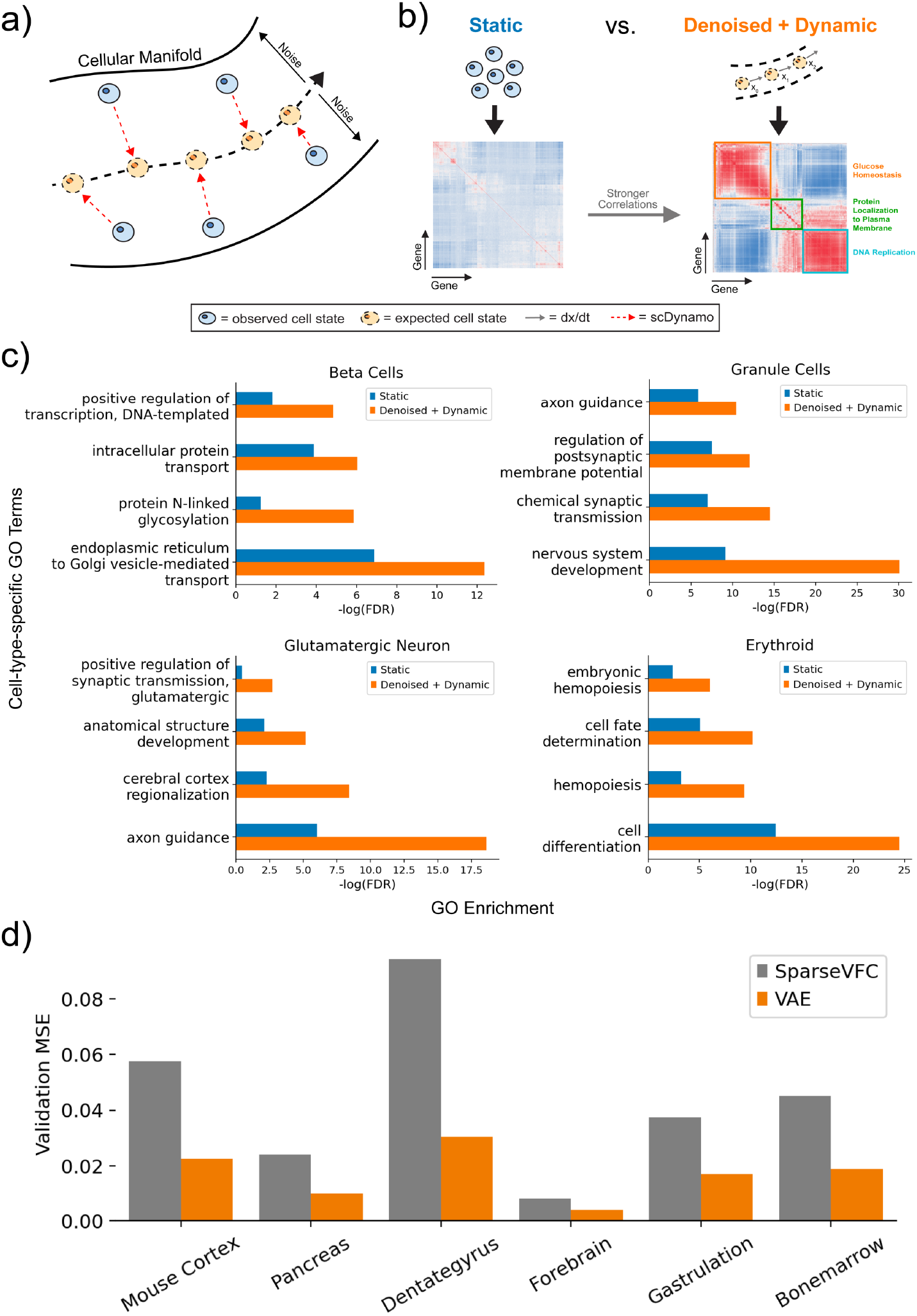
Disentangling Trajectory-specific Gene Co-expression Networks. **a)** Schematic for reducing variability along a developmental trajectory due to sparsity and noise in scRNA-seq experiments with denoising VAEs in DeepVelo. **b)** We used a representative initial condition (e.g., the median expression profile for a cluster of cells) to simulate denoised cell trajectories according to the dynamics learned by DeepVelo. Compared with static cell clusters, the dynamic cells in denoised trajectories have stronger correlations between genes, which leads to better coherence between functional gene modules. **c)** GO enrichment of cell-type-specific terms from the most significant functional gene module. Our method improves upon existing gene co-expression network approaches for cell-type-specific GO term enrichment by at least two orders of magnitude. **d)** Benchmarking with the state-of-the-art vector-field learning method *SparseVFC* shows that our VAE-based framework can improve out-of-sample velocity vector prediction accuracy by at least 50% across the six datasets, indicating that DeepVelo can provide a better representation of the velocity field.

### Comparing DeepVelo with Existing Methods

DeepVelo qualitatively differs from existing ODE-based regulatory networks (*41*). First, explicitly deriving differential equations for biological processes is only feasible when examining small-scale systems (*42*–*45*). In contrast, DeepVelo can capture high-dimensional interactions and can scale to a large number of variables. Second, DeepVelo uses a neural network to learn potentially non-linear gene interactions, which is more suitable for modeling complex biological processes compared with linear ODEs and other kernel-based sparse approximation methods (*46*, *47*). In particular, we compared DeepVelo with the state-of-the-art vector-field learning approach *SparseVFC* (*21*). Benchmarking results showed that our method achieves a reduction of at least 50% in out-of-sample velocity prediction loss across all datasets, indicating that DeepVelo can provide a more accurate representation of velocity vector fields and can compute future cell states with better numerical precision (Fig. 5d). Lastly, many previous ODE-based methods use pseudotime as a substitute for time. In comparison, DeepVelo uses RNA velocity, which reflects developmental time (*20*).

## Discussion

Although many effective tools have been developed for illuminating broad trends in single-cell data, most existing methods view single-cell datasets as a static manifold (*13*, *14*, *8*). In reality, many underlying biological processes captured by single-cell sequencing are dynamical systems, in which individual cells are continuously transitioning from one state to another. Hence, by deriving accurate differential equations that quantify the gene expression dynamics of single cells, we can answer many questions about cell fates and the genetic drivers governing developmental trajectories.

Explicitly deriving differential equations for all gene interactions is a challenging task. Therefore, we tackled the problem with a data-driven approach. We considered each cell in scRNA-seq as an instance sampled from a dynamic system, composed of a state vector (gene expression, 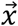) and a velocity vector (RNA velocity, 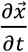). Then, we trained a neural network 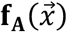 to learn potentially non-linear mappings from the state 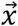 to the velocity 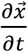 of each cell. With a trained VAE that takes part in the differential equation 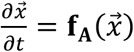, we can integrate the VAE with any black-box ODE solver to compute future (or past) gene expression states.

Overall, our DeepVelo framework learns a continuous vector-field representation of single-cell velocities and allows hypothetical cells to evolve according to the dynamics learned from existing cells in the data. Using the ability to simulate future gene expression trajectories, we devised a metric, the CCI, to quantify the instability of individual cells. By perturbing cell states with high criticality, *in silico* gene perturbation studies can computationally identify key upstream driver genes that determine the fate of cell bifurcations. Lastly, by reversing the developmental time of differentiated cells, retrograde trajectories can deconvolute trajectory-specific gene co-expression networks and discover more coherent cell-type-specific gene modules.

Previous approaches have utilized pseudotime to construct a temporal ordering of cells and pluripotency metrics to measure the differentiation potential of a cell, similar to quantifying the “potential energy” of a Waddington landscape. However, these “potential energy” metrics are limited in describing dynamical systems. Theoretically, the potential energy is converted into conservative forces, where the total work done by a cell becomes independent of the developmental path taken. To accurately measure the magnitude of gene expression changes along developmental paths, we designed the CCI metric, analogous to the “kinetic energy” of a Waddington landscape. In our analysis, we demonstrated that this metric can highlight fixed points in single-cell dynamical systems. Moreover, previous studies have formulated cell fate decisions as high-dimensional critical state transitions (*49*, *50*). Therefore, we hope to bring awareness to the dynamical perspective of single-cell data and advocate for new metrics that quantify the kinetics of single-cell experiments.

More interestingly, it has long been hypothesized that single-cell processes may exhibit properties similar to those of chaotic systems (*51*–*53*). By recovering single-cell gene expression dynamics with DeepVelo, we observed chaotic behaviors in *in silico* gene perturbation studies, in which a small change in the initial gene expression state may result in a large difference in future states, also known as the butterfly effect. Specifically, small perturbations in developmental driver genes of progenitor cells can alter the cell fate at developmental branching points both *in vitro* and *in silico*. If single-cell dynamics exhibit chaotic properties, under the right biological conditions, the chaos can spontaneously evolve into lockstep patterns according to the Kuramoto model of synchronization (*54*). Hence, synchronization models could provide a possible explanation for emerging tissue-level behaviors from single cells. Future works could explore these effects by incorporating the gene interaction dynamics between cells. Currently, DeepVelo only models gene expression dynamics within a single cell. A future direction could expand the state space of DeepVelo and incorporate gene interactions between spatially neighboring cells with spatial transcriptomics (*55*). Another future direction includes incorporating concurrently resolved protein and chromatin accessibilities and their velocities into the dynamical model as a multi-modal representation of the cell state (*56*, *57*)

## Materials and Methods

### Data Collection and Preprocessing

scRNA-seq data (mouse pancreatic endocrinogenesis, dentate gyrus, gastrulation, neocortex, and human forebrain) were downloaded from the NCBI Gene Expression Omnibus repository and the Sequence Read Archive. The raw FASTQ files were preprocessed using *Cellranger* v6.0.1 with default parameters (*58*). As references, we used GRCh38 (2020-A) for human samples and mm10 (2020-A) for mouse samples. After preprocessing, we obtained an mRNA count matrix with rows as cells and columns as genes. Then, using the *Cellranger* intermediate outputs, we computed the amount of spliced and unspliced mRNA transcripts using the *velocyto* package with default parameters (*16*). The output was a loom file with spliced and unspliced mRNA counts for each cell and gene.

For preprocessing and computing RNA velocities, we followed the procedure recommended by *scVelo* (*31*). We selected the top 3,000 highly variable genes with at least 20 mRNA reads, and normalized the mRNA counts within each cell (using the *scv.pp.filter_and_normalize* function in *scVelo*). The normalized mRNA count matrix became the gene expression state matrix. Then, we log-transformed the normalized spliced and unspliced counts for moment calculations. First- and second-order moments were computed using the top 30 principal components and the top 30 nearest neighbors (using the *scv.pp.moments* function in *scVelo*). After recovering the dynamics using the moments, RNA velocity was computed with the generalized dynamical model from the raw normalized reads (using the *scv.tl.velocity* function and *mode* = “*dynamical*” setting in *scVelo*). Only the velocity genes were used as features for the neural ODE. The numbers of cells and available velocity genes for each dataset are shown in the table below.

**Table 1:**
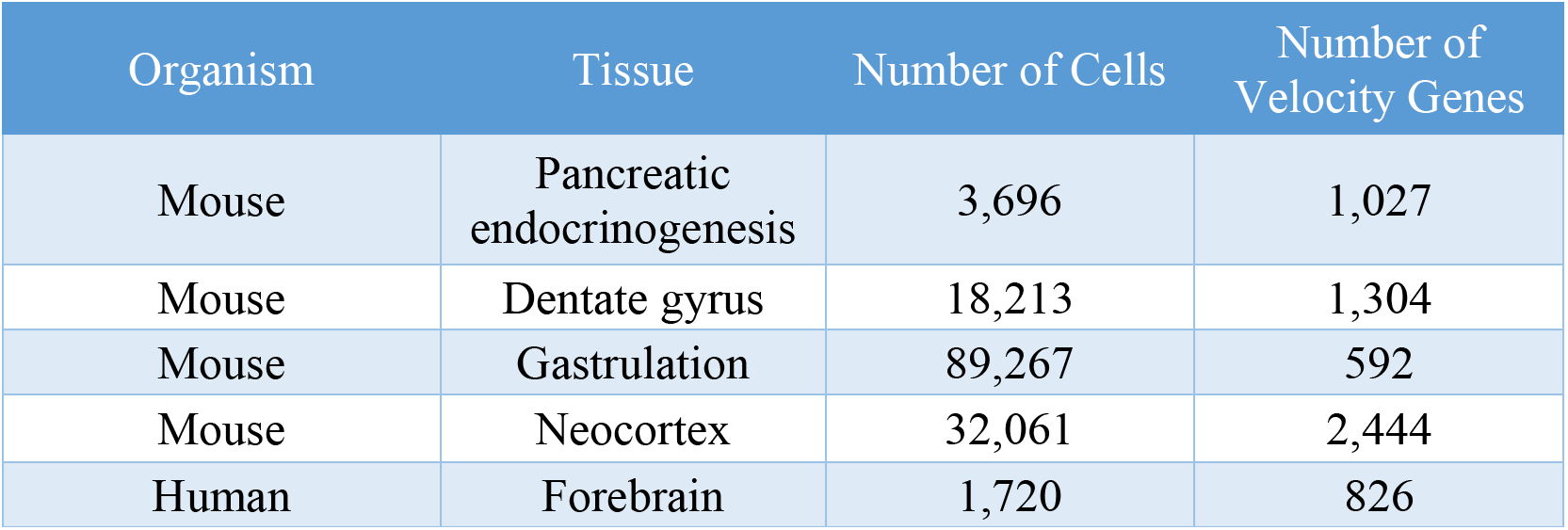
Summary of scRNA-seq datasets used in this study

| Organism | Tissue | Number of Cells | Number of Velocity Genes |
| --- | --- | --- | --- |
| Mouse | Pancreatic endocrinogenesis | 3,696 | 1,027 |
| Mouse | Dentate gyrus | 18,213 | 1,304 |
| Mouse | Gastrulation | 89,267 | 592 |
| Mouse | Neocortex | 32,061 | 2,444 |
| Human | Forebrain | 1,720 | 826 |

### Variational Autoencoder Architecture

High-dimensional single-cell dynamical systems are difficult to model due to high degrees of freedom. For example, the number of features can sometimes be larger than the number of data points. Consequently, gene expression would only vary in a small portion of dimensions. Therefore, modeling the gene expression dynamics of a low-dimensional manifold embedded in high-dimensional data is a challenging task. Fortunately, autoencoders can reduce the dimensionality of the data by introducing an information bottleneck. Accordingly, when used to represent dynamical systems, autoencoders can restrict cell transitions to only movements along the low-dimensional manifold.

A VAE consists of an encoder, which parametrizes Gaussian distributions to be sampled from, and a decoder, which transforms the sampled values into the output. For the encoder and the decoder, four dense layers (size 64 as the intermediate layer and size 16 as the latent layer) with *relu* activation were constructed using the Tensorflow and Keras packages (*59*, *60*). The VAE takes the gene expression state as input, and outputs the RNA velocity. In the VAE, the encoder layers with weights *W_e_* and biases *b_e_* produce the hidden layer *h*(*x*), which parametrizes the location and scale of *i* Gaussian distributions. Then, a sample from each reparametrized Gaussian distribution *z_i_* is used as input for the decoder layer with weights *W_d_* and biases *b_d_*. The architecture can be expressed as:

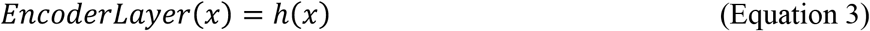

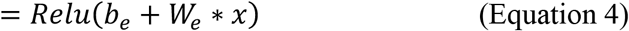

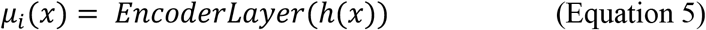

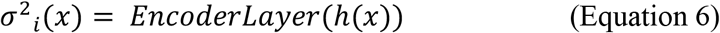

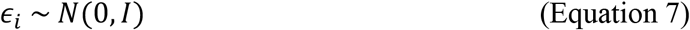

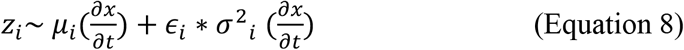

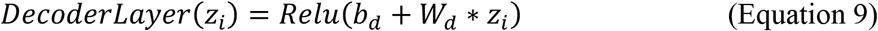

where the *Relu*(*z*) activation function is:

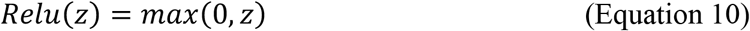

We used the mean squared error as the reconstruction loss and minimized the loss with the Adam optimizer. To prevent overfitting and to encourage a sparse representation of latent embeddings, L1 regularization was added to the activation of all layers with *λ* = 1 × 10^-6^. The evidence lower bound loss function with 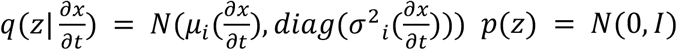, can be described as:

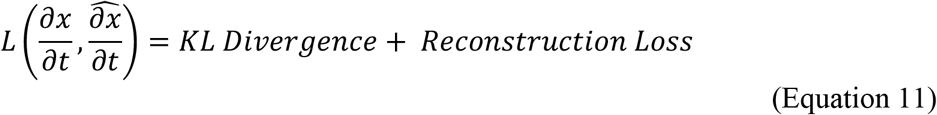

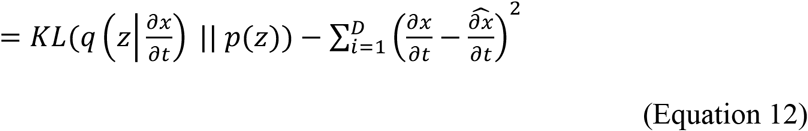

Because the input and output vectors are sparse, a small learning rate of 0.00001 was used with a batch size of 2. Early stopping was added once the validation loss did not improve for three consecutive epochs.

### Initial Value Problems and ODE Solvers for Integration

Our framework can be used to predict gene expression profiles across time. Given *t_0_* and 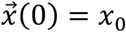, this is an initial value problem with the goal of solving 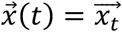 for any *t*:

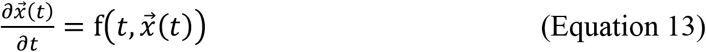

Here, f is only a function of the state 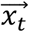 such that 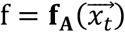. Then the equation becomes:

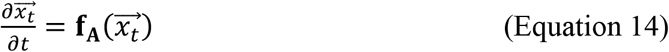

The first-order Euler’s method for finding the state 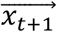 is:

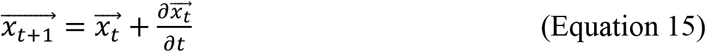

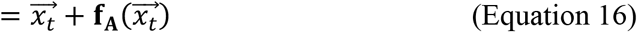

However, we can utilize higher-order ODE solvers from the SciPy package to find a more accurate solution (*61*). The explicit Runge-Kutta method of order 8 (DOP853) was used to obtain the most accurate solutions, but it has a slow runtime. The explicit Runge-Kutta method of order 3 (RK23) can be used to trade off accuracy for a faster runtime.

From the preprocessing steps, gene expression state *x* is the normalized mRNA count, and RNA velocity 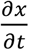 is the rate of change of the normalized mRNA counts. The predicted RNA velocity should be linearly scaled before being used to approximate numerical derivatives with respect to time for integration. We estimated that a velocity scaling factor of 0.65 ± 0.05 worked well in terms of our normalization and integration procedures. Further, to estimate the maximum number of steps to integrate to reach the terminal state of a cell, we computed the 2 standard deviation range for the expression of each gene. Then, we divided the range of each gene by the mean velocity to find the step size of each gene. The maximum step size is defined as the 95^th^–99^th^ percentile of the step sizes. More simply, the maximum step size would allow a 2 standard deviation change in the expression of at least 95% of the genes.

### Addressing Drift Effects

In control theory, using only the previous state and the velocity vectors to predict the next state can result in a phenomenon called “dead reckoning,” where the errors accumulate after each step (*62*). Under our framework, RNA velocity prediction errors could come from many factors. For example, the VAE may not have enough internal (e.g., genes) or external (e.g., environmental) features to accurately predict RNA velocities. The black box integration procedures may also introduce numerical errors. To mitigate this effect, we employed two strategies:

1. Instead of a traditional VAE, we trained a denoising VAE to reduce the variance of the predicted RNA velocity. By adding a small Gaussian noise ϵ (*σ* < 10^4^) to the gene expression input during training, we could increase the generalizability of the input space and improve extrapolations to out-of-sample cell states.

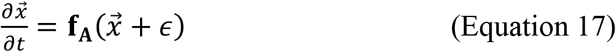
2. As we integrated the VAE over time, we found reference cells in the data manifold every few steps and continued integration from the reference cell, as a form of a high-gain Kalman filter. We designated the intermediate step size as a hyperparameter relative to the step size. For example, after integrating for five intermediate steps, we projected the predicted (or extrapolated) gene expression state to the original dataset using the top 30 principal components. Then, we identified the K-nearest neighbors (*K* = 30) within the PCA embeddings. The reference cell is defined as the median expression profile among those K-nearest neighbor cells from the dataset, and ODE integration continued from this reference cell. This allowed our prediction to adhere closely to the data manifold and further reduced the degree-of-freedom. Consequently, finding reference cells in the data also constructed boundary conditions when integrating a dynamical system. For example, once the extrapolated state went beyond the cellular manifold, there were no cells in the data to serve as a reference, but the nearest neighboring cells from the dataset could still construct a reference cell from where integration could continue.

### Measuring Instability with the Cell Criticality Index

By solving for the developmental path of a single cell, we can measure the amount of gene expression change along a trajectory, rather than comparing only the difference between the start and end states. Previously, StemID used the entropy of the gene expression distribution to heuristically identify stems cells in single-cell transcriptome data, where pluripotent cells tend to have a more uniform gene expression distribution with a higher entropy and differentiated cells tend to have a lower entropy (*63*). If 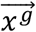 denotes the expression state of the genes *g*, then the StemID of the gene expression state is defined as:

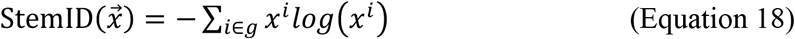

We reasoned that a change in the gene expression distribution (e.g., from high to low entropy) can be captured using the relative entropy (or the KL-divergence). Based on this idea, we devised a measure to quantify the capacity for any cell to undergo a gene expression change in the dynamical system. The CCI is calculated as the cumulative information change, or the cumulative KL-divergence, between gene expression distributions at each step in the developmental trajectory. Different from StemID, the CCI can quantify the gene expression change of a cell regardless of the pluripotency. As an analogy, StemID measures the “potential energy” of a cell’s ability to differentiate, whereas the CCI measures the “kinetic energy” of a cell’s ability to change. If 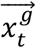 denotes the expression state of the genes *g* at time *t*, then the cumulative KL-divergence for *T* = 35 steps can be defined as:

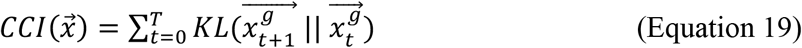

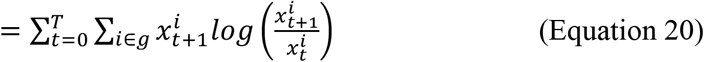

#### Sampling Out-of-sample Cells and Simulating Perturbations

To simulate out-of-sample initial gene expression states, we computed the median expression profile of a certain cell type (e.g., pre-endocrine progenitors in pancreatic endocrinogenesis) and added Laplace-distributed (two-sided) noise using the variance of each gene within that cell type to randomly increase or decrease gene expression.

For the perturbations, exponentially distributed (one-sided) noise was added only to the top 100 differentially expressed genes for the randomly sampled cells to specifically increase the expression of the top differentially expressed genes. Terminal cell identity was determined by projecting the data onto the top 30 principal components and by using K-nearest neighbor classification (with *K* = 30). With the *scVelo* package, the dynamical mode estimates a variance for each gene over all cells, whereas the stochastic mode estimates a variance for each cell. Note that to model stochasticity in the stochastic mode, our framework could be easily adapted to also learn the variance of the velocity vectors (as neural stochastic ODEs).

### *In Silico* Perturbation Studies

We divided the *in silico* perturbation study into three steps:

1. A sample of initial gene expression states (*n* = 1,000) was randomly generated. First, we solved the random initial gene expression states over time to establish a developmental baseline. Specifically, we aimed to observe the natural proportion of terminal cell types that could arise from the dynamical system without any intervention.
2. Then, we identified differentially expressed genes in the initial gene expression states that correlate with development into a particular terminal cell type later in time. Differential gene expression was performed using the *scanpy* package with the Wilcoxon test and Bonferroni corrections (*64*).
3. Lastly, we perturbed only the differentially expressed genes in another set of randomly sampled initial gene expression states to test whether the perturbation increases the proportion of cells for the terminal cell type.

### Retrograde Trajectory Simulation

Similar to the *in silico* perturbation studies, we computed the median expression profile of a terminal cell type (e.g., beta cells, granule cells, glutamatergic neurons, or erythroid cells) in each scRNA-seq experiment (mouse pancreatic endocrinogenesis, dentate gyrus, human forebrain, and mouse gastrulation) as the representative initial condition. A set of cells (*n* = 50) was sampled from each representative initial condition by adding Laplace distributed noise using the variance of gene expression of the terminal cell type. The retrograde trajectory for each cell was simulated by subtracting the predicted RNA velocities from the gene expression state during integration:

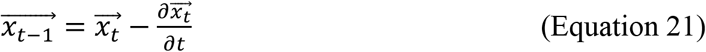

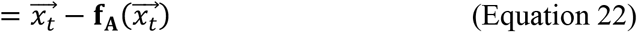

After integrating for 15 discrete steps each with 5 intermediate steps, a gene correlation matrix of the cells in retrograde trajectories was calculated.

### Gene Ontology Enrichment Analysis

Hierarchical biclustering was performed on the co-expression matrices, and three gene clusters were identified from each co-expression matrix, representing three functional modules. We performed GO enrichment analysis on each functional module using GOATOOLS with Fisher’s exact test (*65*). Further, we calculated the Benjamin-Hochberg false discovery rates to correct for multiple testing. To compare between two co-expression matrices, we considered the most significant enrichment out of the three clusters for each GO term. In Fig. 3a and Fig. 5a, the most significantly enriched GO terms associated with biological processes are listed next to each gene cluster.

## Author contributions

Conceptualization: ZC

Methodology: WK, ZC

Investigation: ZC, WK, MB, JZ

Visualization: ZC

Supervision: MB, JZ

Writing—original draft: ZC

Writing—review & editing: MB, JZ

## Data and materials availability

The mouse pancreatic endocrinogenesis, dentate gyrus, neocortex, gastrulation, and human forebrain datasets used for this study can be found in the NCBI Gene Expression Omnibus repository with accession numbers GSE132188, GSE95753, GSE153164, and GSE87038, and in the Sequence Read Archive under accession code SRP129388, respectively. All source code to reproduce this study can be found on GitHub at https://github.com/gersteinlab/DeepVelo.

