## Supplement for "DeepVelo: Single-cell Transcriptomic Deep Velocity Field Learning with Neural Ordinary Differential Equations"

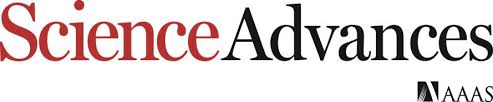

Supplementary Materials for

**DeepVelo: Single-cell Transcriptomic Deep Velocity Field Learning with Neural Ordinary Differential Equations**

Zhanlin Chen, William C. King, Mark Gerstein, Jing Zhang

**This PDF file includes:**

Figs. S1 to S5

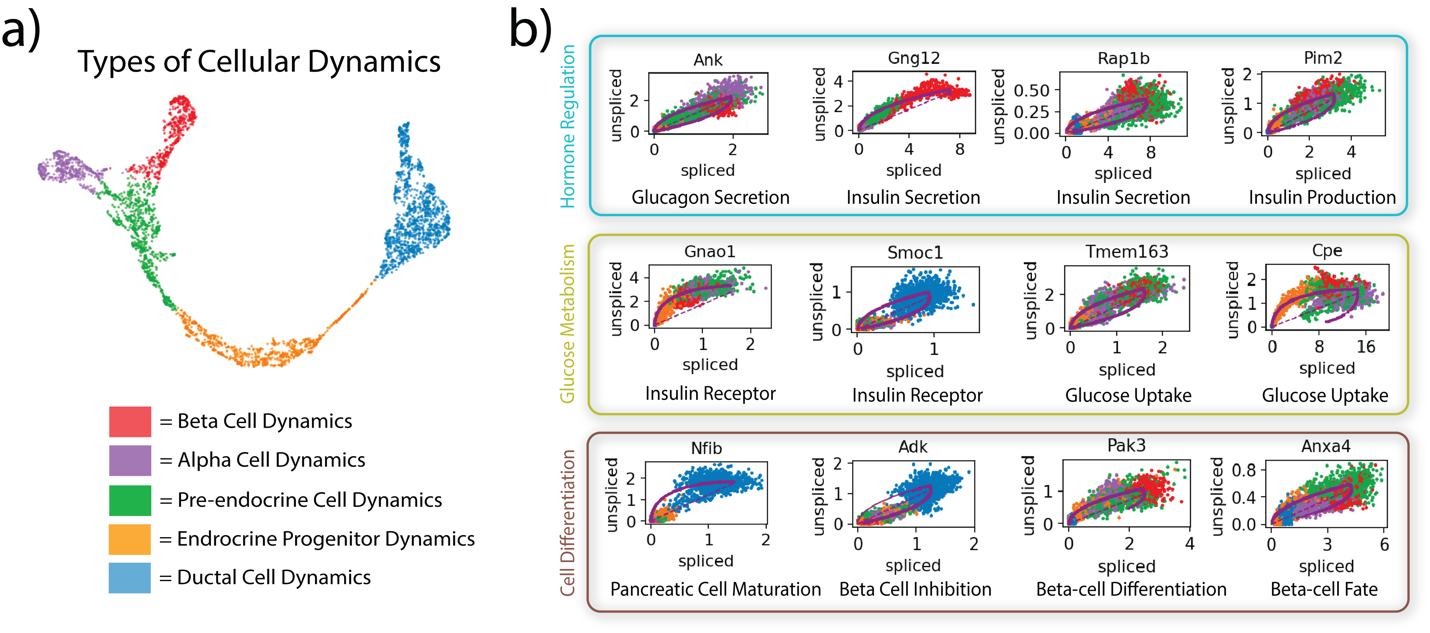

Fig. S1. Pancreatic Endocrinogenesis Differential Kinetics. a) Clustering the latent layer embeddings from our framework can find distinct types of cellular dynamics in pancreatic endocrinogenesis. b) Differential kinetic testing between different types of cellular dynamics reveals genes crucial to hormone regulation, glucose metabolism, and cell differentiation.

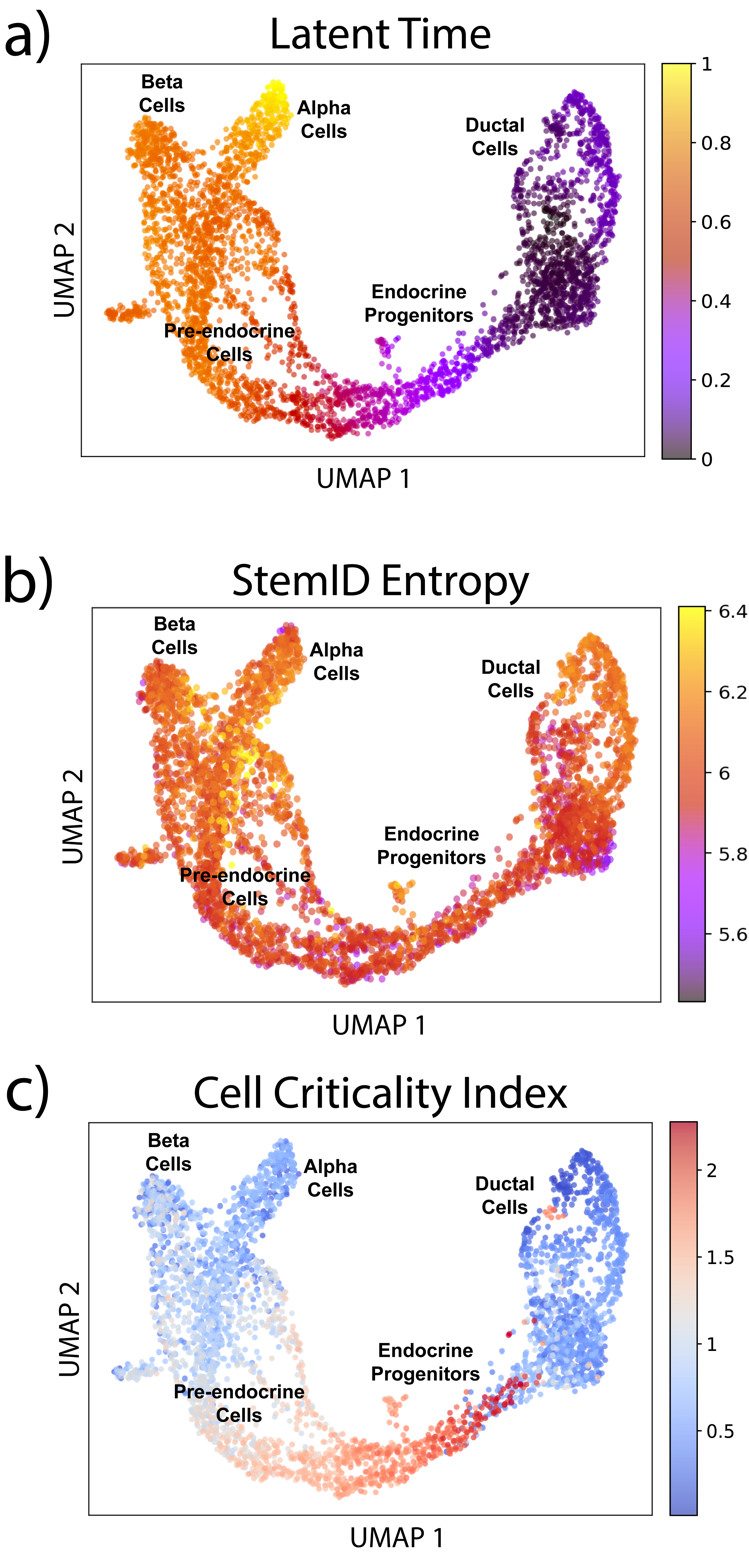

Fig. S2.

**Comparing Velocity-based Latent Time, StemID Entropy, and Cell Criticality Index**. In pancreatic endocrinogenesis, Velocity-based Latent Time provides a pseudotime temporal ordering of cells. StemID Entropy is designed to order the cells by pluripotency potential, but it fails here because the cells have similar amounts of uniformity in gene expression. Lastly, Cell Criticality Index highlights fixed points in the developmental landscape.

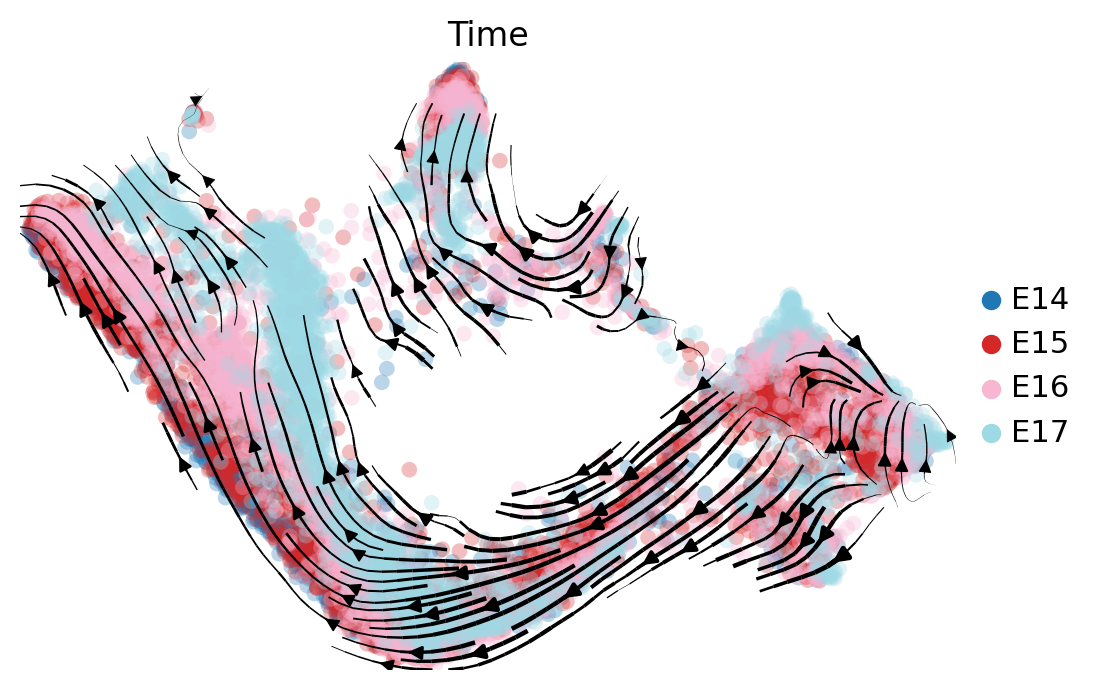

Fig. S3.

**Batch Correction for Developing Mouse Neocortex.** After filtering for the top 5,000 highly variable genes and normalizing within each time group, the “sc.pp.combat” function from *scanpy* was used to correct for batch effects. Further, the velocities were computed within each time group.

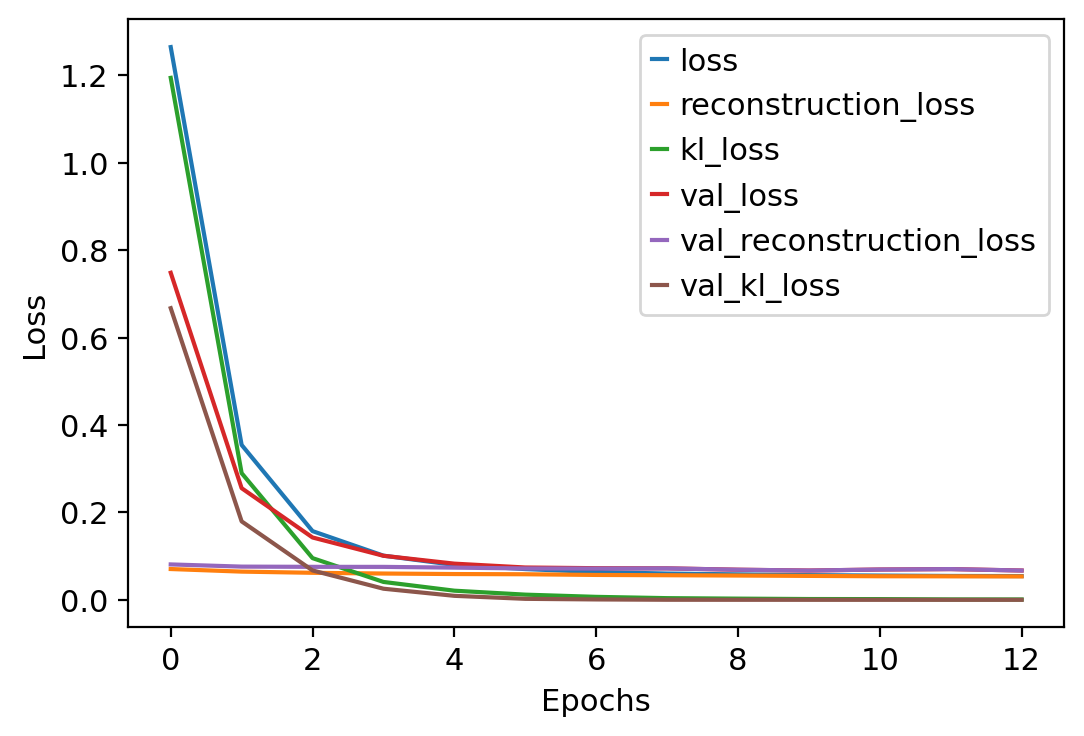

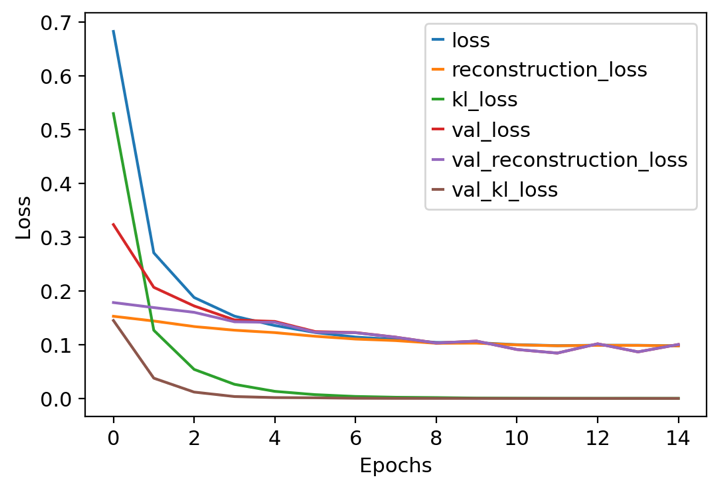

Fig. S4.

**Variational Autoencoder Training.** The training and validation loss (reconstruction, KL-divergence, and reconstruction with KL-divergence) of the variational autoencoder during training (left – pancreatic endocrinogenesis, right – dentate gyrus).

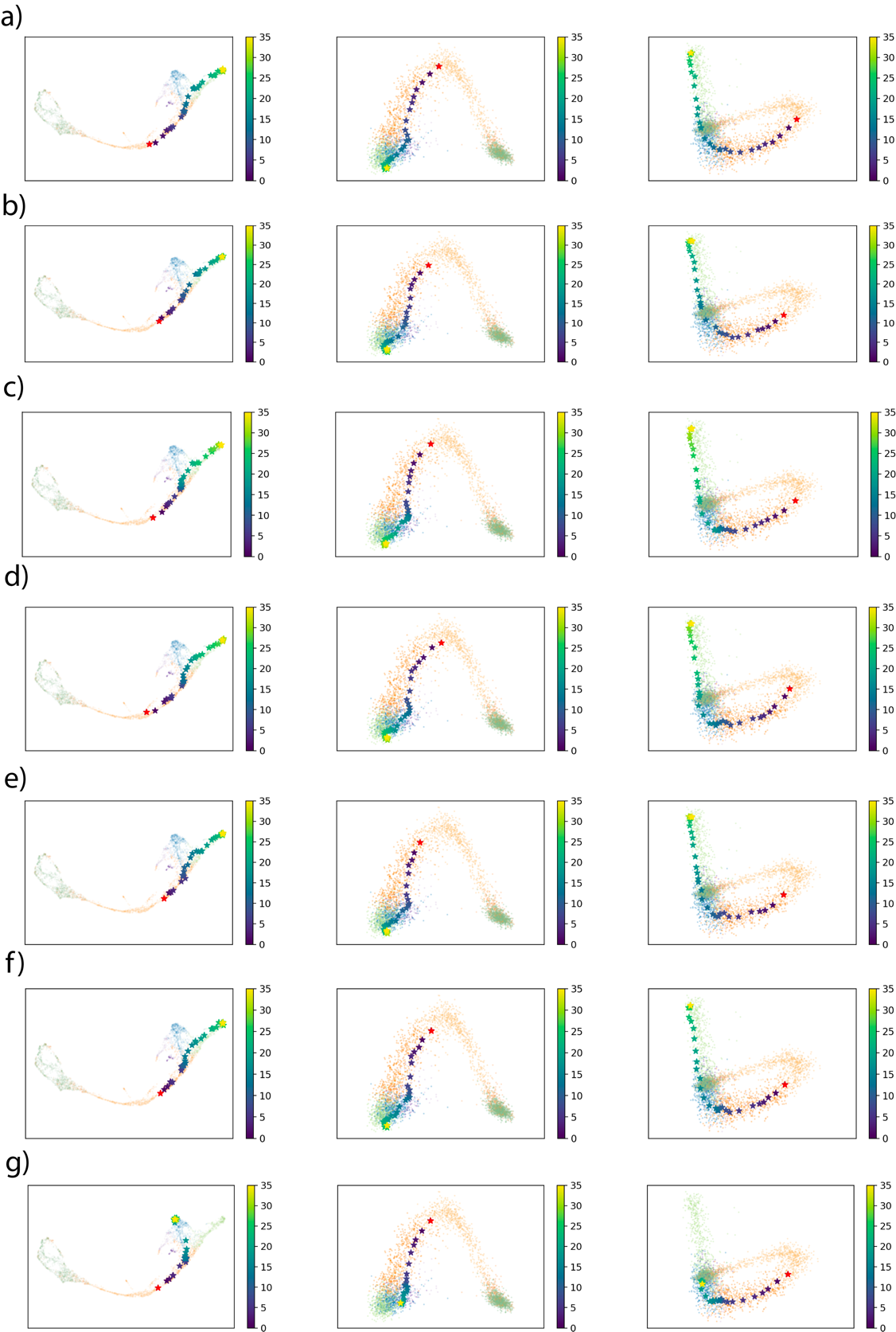

Fig. S5.

**Example Pancreatic Endocrinogenesis Trajectories.** **a-f)** Endocrine progenitor developing into pre-endocrine cells, then differentiating into beta cells. **g)** Alternative trajectory of progenitor cells developing into alpha cells.
